# Sound-evoked orofacial motion only affects a fraction of visually responsive V1 neurons in a multisensory detection task with delayed reward delivery

**DOI:** 10.64898/2026.06.05.730357

**Authors:** Medina Husić, Reinder Dorman, Jeannette A.M. Lorteije, Umberto Olcese, Cyriel M.A. Pennartz

## Abstract

Auditory stimuli have been known to elicit neural responses in visual cortical areas, but such responses are partly associated with sound-evoked orofacial movements. However, it remains unclear to what extent neural correlates of movement can be dissociated from non-motor correlates. Here we developed a paradigm with a forced delayed-response mechanism by means of a moving reward apparatus in order to test whether this results in low amounts of movement-related neural activity shortly after stimulus onset. Animals performed a multisensory detection task, while we recorded neural activity from V1 and measured orofacial motion. During the delay, when only the stimulus was present, orofacial motion was relatively low. Reward-related motion increased after the delay, with a concomitant increase in motor-related spiking activity. Visual stimulation did neither elicit considerable motion nor corresponding neural responses in the delay period. Conversely, salient auditory stimuli led to modest, but increased orofacial motion and related neuronal activity during the delay. We identified a subpopulation of visually responsive V1 neurons that did not show correlations with orofacial movement, with only the stimulus being represented. These results present a forced delayed-response window as a method to help disentangle motor-related activity from visual (but not auditory) processing and show that motor-related activity only affects a fraction of visually selective neurons.

## Introduction

One of the main goals in neuroscience is to understand the neurobiological underpinnings of complex behaviors. In rodents, this is accomplished by combining neurophysiological measurements, such as *in vivo* electrophysiology, with complex behavioral tasks. In these tasks, mice are typically trained to respond to stimuli, allowing the researcher to establish correlational and/or causal connections between stimulus and response. However, it has become apparent that it is important to account for body-movement confounds in neural correlates^1–4^. Movement-related signals may temporally and spatially overlap with processes considered sensory or cognitive. Tasks are usually designed to separate such processes from motor execution by means of a learned delay to respond behaviorally after stimulus presentation^5–8^. Nevertheless, detectable preparatory activity can remain present during the delay and stimulus presentation^9^. This activity prior to self-initiated movement is present cortex-wide^10^. This complicates the separation between neural correlates of motor-related and sensory/cognitive activity.

Not only task-related, but also spontaneous, uninstructed movements can confound the interpretation of results on neural encoding of cognitive processes^2–7,11,12^, without requiring primary motor cortex (M1)^13^. These signals are widespread throughout the dorsal cortex, including primary sensory areas, and are linked to anticipatory movements and premotor planning ^4–6,10^. For example, a considerable portion of the variance in neural activity in the primary visual cortex (V1) during awake, passive viewing of visual stimuli can be explained solely by spontaneous movements^4^. Likewise, auditory responses in V1 have been found to be contaminated with or evoked by task-unrelated orofacial movements^2,3,14^. Sound, orofacial movement and locomotion may even enhance neural encoding of visual stimuli in V1^15,16^.

Contextual and Pavlovian influences also play a role in shaping V1 activity, further complicating the disentanglement of sensory, cognitive, and motor signals. The addition of Pavlovian cues in a behavioral task can unwillingly enhance instrumental behavioral responses through Pavlovian instrumental transfer^17^. These can in turn be accompanied by enhanced neuronal activity. Moreover, Pavlovian cues can affect non-instrumental behaviors and neural processing in sensory areas. For example, stimulus-reward associations and reward expectation alter stimulus-specific neuronal populations and sharpen V1 representation of retinotopic space^18–20^. Thus, Pavlovian task variables such as reward-predicting detection cues or reward delivery devices can affect uninstructed or instrumental movement, neural activity evoked by conditioned stimuli or a combination thereof. Task design can therefore greatly influence motor confounds in neural activity, but how to minimize these remains an open question.

Here, we investigated whether a forced, delayed introduction of the reward apparatus (instead of a learned delayed response) is accompanied by relatively minor levels of motor correlates in sensory responses of V1 neurons, bearing in mind that the proximity of the reward apparatus may elicit considerable motor activity even before reward is delivered. We recorded V1 single units in mice trained on a multisensory 2-alternative-forced-choice stimulus detection task with a forced delay between stimulus onset and behavioral response to acquire reward. Mice had to indicate the presence or absence of an audiovisual stimulus, but could do so only after a short delay relative to stimulus onset. The end of the delay period was marked by a dual-lickspout moving into position to be reachable for the mouse. Lickspout arrival induced orofacial movement, possibly enhanced through the Pavlovian effects already referred to. Salient auditory, but not visual, stimulation elicited considerable orofacial movement before instrumental lick responses were made. Thus, the stimulus window does not remain free of orofacial movement for all modalities. These movements were strongly correlated with single unit activity in V1, but only in a subpopulation of neurons. A Random Forest encoding model showed that a sizable subpopulation of visually selective units remained largely free of motor confounds in the initial visual response period. Contrarily, auditory-evoked responses in V1 were largely explained by orofacial movements. Thus, in a task design with a forced response-delay relative to the stimulus, orofacial motor responses and their V1 correlates are limited during visual, but not auditory stimuli.

## Methods

### Animals

Five adult male C57/BL6j mice (Envigo; 2 PVCre and 3 PVCre-TdTomato), at least 8 weeks at the start of the experiment, were group housed in a climate-controlled room (21 ± 2 °C and 55 ± 15% humidity). Age during recordings ranged from 80 to 315 days. The animals were put on a reverse light cycle and all experimental procedures were conducted in the dark phase (08:00 – 20:00 hrs). Food was available *ad libitum*. Water intake was restricted to maintain at least 85% of the free-feeding weight. Typically, daily water intake was obtained during training (1-2 mL), but was supplemented with Hydrogel™ if the minimum intake (0.025 ml/g body weight) was not met. If signs of dehydration occurred, animals were taken off water-restriction until fully recovered. All experimental procedures were approved by the Dutch Commission for Animal Experiments and by the Animal Welfare Body of the University of Amsterdam under protocol AVD1110020172385.

### Behaviour

Mice were trained on a multisensory two-alternative-forced-choice stimulus detection task with a delay between stimulus and instrumental response to acquire reward. There were four trial types: Visual-only (V), Auditory-only (A), multisensory (M), and no-stimulus (probe, P) trials. Mice had to indicate whether they had perceived a stimulus or not by licking either the right or left side of a movable Y-shaped two-sided lickspout, respectively. Upon stimulus presentation, after a random delay (0.5-1.5 s), a servomotor moved the lickspout towards the snout, which allowed the animal to respond. The period between stimulus onset to servo onset was defined as the stimulus-only epoch (0-0.5 s), and the period in which the servo could arrive (0.5 – 1.5 s) was defined as the lickspout-arrival epoch. This was followed by a response/reward epoch which lasted until the end of the stimulus presentation (4 s). The servo moved away from the mouse after reward consumption (2 s after correct lick) or at the end of the stimulus period (if no lick had been forthcoming).

#### Training

Before commencement of behavioural training, animals were gradually habituated to experimenters, the setup and head-fixation. The training protocol started with a conditioning stage, in which we randomly presented stimuli (A or V) and probe trials. The servo moved the lickspout to the mouse after a random delay between 500 and 1500 ms, after which a reward was given without a requirement for an operant response. The next trial started after a random 3-7s inter-trial interval (ITI). Once mice licked consistently upon stimulus detection, they moved to the active stage of training where they received no rewards while being passive; only correct responses were now rewarded to promote active licking. Incorrect responses were followed by the servo moving away from the mouse immediately and a 4 s time-out. The active stage consisted of visual-only and auditory-only blocks of 5-50 trials, which decreased in size as training progressed. Once mice were performing well, we added the multisensory condition (i.e. with a simultaneous auditory and visual stimulus) and presented the auditory, visual and multisensory stimuli conditions pseudo-randomly. If the mouse was strongly biased towards one side of the lickspout, we repeated the stimulus condition resulting in repetitive errors as a countermeasure. Next, contrasts of the stimuli were lowered to eventually train mice on the full task.

#### Full task

Contrasts of stimuli were lowered individually per animal to detection threshold (50% correct detection or hit rate). The full task consisted of three contrasts per stimulus type: high (i.e. maximum, approaching 100% detection rate), mid (i.e. near threshold, 50-75% detection-rate) and low (i.e. sub-threshold, below 50% detection rate). Lower contrasts would elicit *unperceived* responses, i.e., licks to the left. To account for those and get a similar amount of responses to either side of the lickspout, we lowered the amount of probe trials to 35-45% of the total trial count. This ratio was estimated based on the individual animal’s amount of *unperceived* responses in the latest training sessions. In the full task the servo moved the lick-detector to the animal after a random 0.5-1.5 s delay.

#### Apparatus

Mice were head-fixed in a sound-attenuated box. The behavioral task was custom-written in OCTAVE (GNU Octave 4.0), using Psychtoolbox3, and ran on a PC running Ubuntu (20.04.1). Visual stimuli were presented at 60 fps on a 21 inch monitor at 23 cm distance from the mouse. Auditory stimuli were high-pass filtered (Beyma F100, Crossover Frequency 5-7 kHz) and delivered through two bullet tweeters (300 Watt) positioned to the right and left of the screen at 35 cm distance from the mouse. During training, licks were registered using custom-made, capacitance-based dual-lickspouts controlled by a microcontroller (Arduino UNO) running the Arduino capacitive sensor libraries (https://docs.arduino.cc/libraries/capacitivesensor/). During recording sessions we used a junction-potential based lick detector^21^ feeding into an analog-digital converter (16 bit ADC and PGA, ADS1115, Texas Instruments, Dallas) and an Arduino UNO running standard Arduino libraries (https://docs.arduino.cc/libraries/). Liquid reward (babymilk; Kruidvat infant formula; 2-5 µl) was dispensed using solenoid pinch valves (Biochem Fluidics) and gravitational force.

#### Stimuli

Visual stimuli were full-screen moving gratings with a spatial frequency of 0.05 cycles per degree, moving at 1.5 Hz with 60 fps. We presented gratings with three different orientations: 0, 120 and 240 degrees. The contrasts used during recordings were adapted to the detection threshold of each individual mouse. During the ITI we showed a grey screen. To match visual stimulation, the pure-tone auditory stimulation was modulated at a frequency of 1 Hz. During training the base frequency was 15 kHz and the depth of modulation was 1 kHz, leading to fluctuations that ranged from 14 kHz - 16 kHz. The base frequency used during recording sessions was adapted to each individual mouse to elicit optimal auditory responses at a maximum volume of 85 dB (which is unlikely to evoke an auditory startle response ^22,23^). During probe insertion in the auditory cortex, we played a pure tone and changed the frequency in steps of 1 kHz until there was visible activity in raw electrophysiological signals (i.e. spiking activity or event-related potential).

#### Video acquisition

Orofacial movements were monitored with a monochrome camera (DMK 22BUC03, The Imaging Source) and zoom lens (50 mm, F/2.8 2/3-inch 10 MP, Navitar) at 60 fps. The camera was positioned at 36 cm distance from the mouse and we used infrared light (850 nm) for illumination. Frames were recorded with a frame grabber for the OpenEphys plugin GUI. Video frames were either 640×480 or 744×480 pixels in size.

### Surgical procedures

#### Head bar implantation

At the start of the experiment (before behavioral training), mice were implanted with a custom-made titanium head-bar. Carprofen (2-5 mg/kg) was injected subcutaneously before surgery as preoperative analgesia. Animals were anaesthetized with isoflurane (4-5% induction; 1-2% maintenance). Temperature was maintained at 36-37 °C with a heating pad. Artificial eye drops were administered and every hour we injected saline subcutaneously to prevent dehydration. The head was shaved and disinfected with an iodide solution (Betadine) and 70% ethanol. We made an incision along the AP-axis, on which we applied 1-2 sprays of Lidocaine (Xylocaine^R^, 100 mg/ml). We removed the periosteum with cotton swabs and cleaned the skull with H_2_O_2_ and 70% ethanol. The temporal muscles were slightly retracted, over which the skin was folded to close the incision. The skin was sealed with tissue glue (Vetbond™) to prevent infection and regrowth. The head-bar was stabilized to the skull with cyanoacrylate glue (Loctite 401, Henkel) and secured with dental cement (C&B Superbond, Sun Medical), while making sure all gaps between the head-bar and skull were filled and there were no sharp edges. The skull was covered with a thin layer of cyanoacrylate glue and silicon adhesive (Kwik-Sil™). The silicone cap was covered with a small metal plate, which was glued to the head-bar and only removed for electrophysiological recordings. Water restriction and training started after a one-week recovery.

#### Craniotomies

Once fully trained the mice were habituated to the experimental setup for a week, after which the mice underwent craniotomy surgery. The procedures regarding anesthesia and analgesia were identical to the procedures described for head-bar implantation. We drilled a small craniotomy (maximal diameter about 200 to 500 μm), measured from bregma, to access left V1 [3.5-3.8 mm AP; ± 2.1-2.5 mm ML]. Craniotomies were kept moist with saline during surgery and covered with silicon (Kwik-Sil™) and a metal plate to protect cortical tissue. The recordings took place after a recovery period of at least 24 hours.

### Electrophysiology

We used 1449 µm 64-channel silicon probes with 23 µm between electrode sites (Neuronexus, Ann Arbor, MI – A1x64-Poly2-6mm-23s-160). We lowered the probes slowly (approximating <0.1mm per sec) to penetrate all cortical layers (800 µm from cortical surface) with manually operated micromanipulators (M3301, World Precision Instruments). The probes were allowed to stabilize for approximately 10 minutes before the start of recording. Signals were acquired with an OpenEphys acquisition system^24^, sampling continuously at 30 kHz and pre-amplified and filtered between 0.1 and 20 kHz with an amplifier board (RHD2164, Intan Technologies). The onset and offset of each stimulus- and servo-event was timestamped by the OpenEphys acquisition system via TTL pulses using an Arduino. The lick timestamps were synced in the behavioral script.

### Histology

At the end of the recordings, mice were intraperitoneally injected with an overdose of Euthasol and transcardially perfused with 4% paraformaldehyde (PFA) in buffered saline. The brains were extracted and kept in 4% PFA for at least 48 hours before sectioning. We sliced coronal sections (50 µm) with a vibratome, which we stained with DAPI (Sigma D9542, 5μl) and cover-slipped with Mowiol (Sigma 81381). We imaged the sections to verify probe placement (Leica MM AF.16).

### Analyses

Unless otherwise stated, all data were analyzed using custom-written Python scripts (Python 3.x). We included 14 sessions from 5 animals (3 ± 0.89 sessions per animal; mean ± SD) in the analysis.

#### Neural data processing

Data was preprocessed by common-average referencing across all active channels. Raw signals were filtered between 600 and 6000 Hz for spike detection. We spike-sorted using Klusta, the Phy Gui ^25^ and Spike Interface software^26^. During manual curation, clusters were included as single units based on the isolation distance (>10) and inter-spike-interval violation (<0.05% within 2 ms ^3,27–29^). To construct peristimulus time histograms spikes were binned in 1 ms time intervals. Bin counts were averaged across trials and smoothed with a gaussian window with a 10 ms standard deviation.

#### Orofacial movement

We used FaceMap^4^ to extract orofacial movements from video data. ROIs containing the moveable lick-detector were manually selected and excluded from analysis. The videos were downsampled by a factor of four. We took the luminance difference between every consecutive frame, and applied singular value decomposition (SVD) on a frame-by-frame basis. This resulted in 500 components explaining orofacial motion across the entire video. We took the absolute sum of these components to calculate the videomotion energy (videoME).

#### Predicting spike trains with a Random Forest model

To estimate how much variance in neuronal activity stimuli and movement explain across time, we predicted spikes by fitting a Random Forest model with 100 trees and 5-fold cross-validation with randomized folds, repeated five times^30–32^. We predicted binned spike counts (100 ms) of single neurons in single trials (-1000 to 4000 ms relative to stimulus onset). We included the visual, auditory, motor, and lickspout-related predictors. The visual and auditory predictors indicated the presence of the stimuli and were expanded in 500 ms bins, creating eight predictors per modality, and also specified the corresponding stimulus intensity (for auditory; three levels) and contrast (for visual; three levels). The motor-related predictors included videoME, lick count to left and lick count to right. The movement of the lickspout was indicated by two predictors: one for moving towards the snout and one for moving away from the snout. Together, we created a 26 (predictors) by 50 (time bins) predictor-matrix for each trial. Encoding quality per time epoch of interest (stimulus-only, lick-detector arrival, or lick/reward epoch) was assessed with the Poisson pseudo-R^2^ score (PR2) averaged over folds and repeats. The PR2 value is a measure of goodness of fit of non-linear regression models and takes a value of 1 when the model perfectly describes the data and a value of 0 when the tested model describes the data similarly to the null model^33,34^.

#### Statistics

Repeated measures ANOVAs and relevant post-hoc t-tests were Bonferroni-corrected and applied where mentioned in the text. A Generalized Linear Model (GLM) was applied to the spike count data, where an ANOVA is not applicable. The GLM we performed in *R*, using the *glm* function of the *lme4* package. The formula used was ‘*difference in spike count ∼ epoch’*, with a binomial Link function to arrive at predicted differences in spike count between epochs. Additionally, the GLM formula ‘*difference in PR2 value ∼ epoch + visually selectivity*’, with a Gaussian link function, was used to determine whether there was a difference in encoding quality between visually selective units and the general population. Pearson correlations were computed to describe correlations between neuronal firing rates and motion energy, and significantly correlated cells were qualified based on Bonferroni-corrected circular shuffle permutations (10.000 iterations), with a threshold of p=0.05. Here, a GLM was used to assess the difference in correlations over time between the two subpopulations of neurons (i.e. visually and auditory selective subpopulations). The formula used was ‘*difference in correlation of firing rates and motionME ∼ neuron subpopulation + neuron ID*’.

## Results

We will first address the question whether a delay in instrumental responding upon stimulus presentation, forced by means of a movable lickspout, is accompanied by limited motor confounds in perceptual neural correlates can be limited Next we will focus on the extent to which sensory responses of V1 neurons can be isolated from motor correlates. We trained five mice on a multisensory two-alternative-forced-choice stimulus detection task (Fig. 1a, b). Four trial types were used: auditory (A), visual (V), multisensory (M), and no stimulus (probe, P). Mice had to indicate the presence or absence of a visual and/or auditory stimulus by licking the right or left side of a dual-lickspout, respectively. Upon stimulus presentation we forced a delay in behavioral responding by moving the lickspout towards the snout of the mouse 0.5-1.5 s after stimulus onset. This resulted in a stimulus-only epoch (0-0.5 s after stimulus onset) wherein mice were or were not presented a stimulus but could not make an instrumental response as the lickspout was out of reach (Fig. 1c). The stimulus-only epoch was followed by the lickspout-arrival epoch during which the lickspout started moving at a variable time within a 0.5-1.5 s time-window. Once the lickspout was in reach of the mouse, the response/reward epoch started (lasting until 4 s after stimulus onset in the case of no response), and mice were rewarded immediately after a correct response. Once the trial ended, the lickspout moved away from the mouse, which was followed by an inter-trial-interval (ITI, 3-7s) marked by a grey screen and no sound. We recorded orofacial movements during task performance and extracted the motion energy per frame using Facemap^4^ (Fig. 1d).

**Figure 1.**
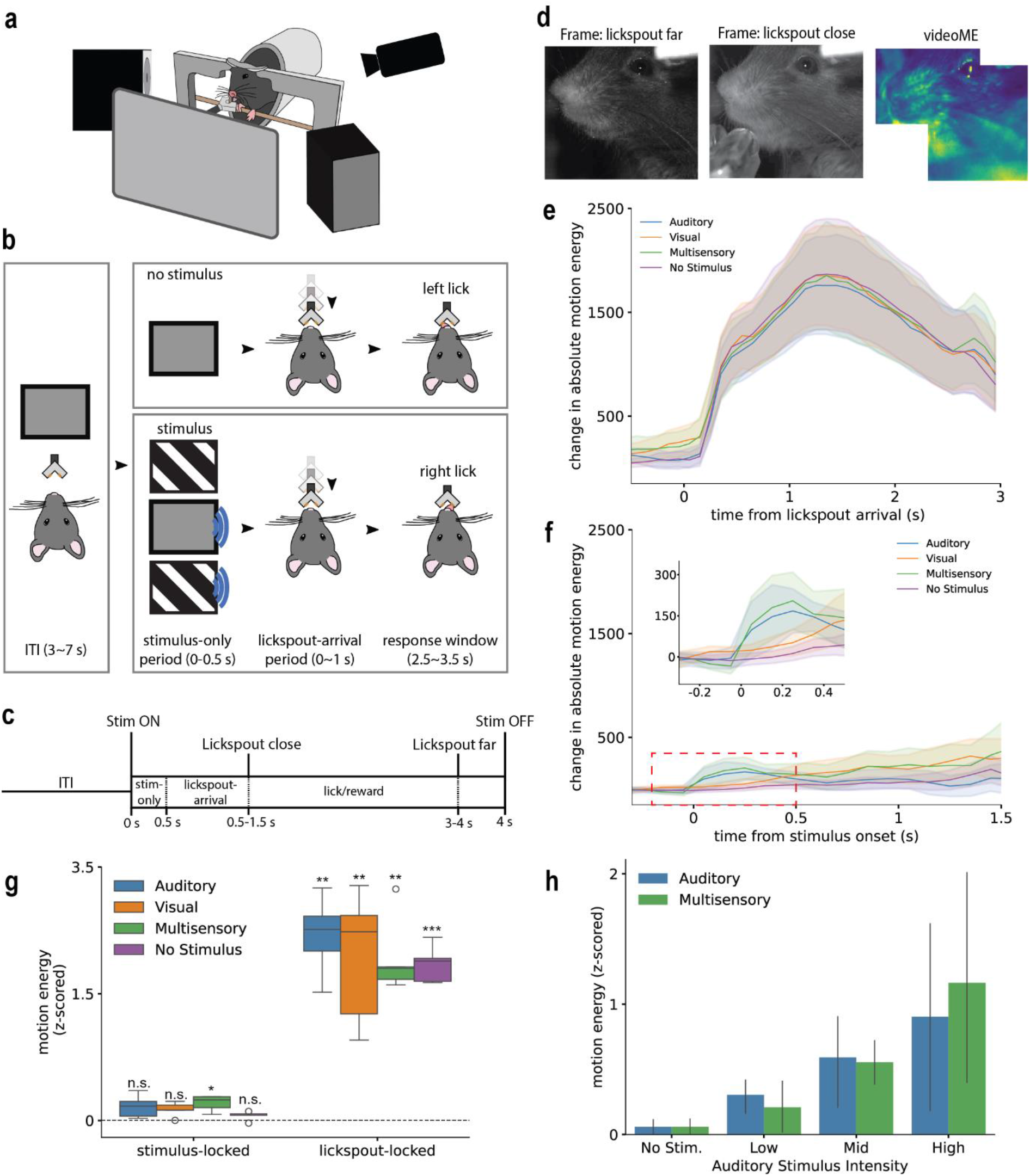
Task-induced increase in motion energy. **a**) Schematic illustration of the behavioral setup for head-fixed mice. **b**) The multisensory detection task. **c**) Timeline for a single trial. Note the lickspout arrives randomly between 0.5 to 1.5 s post-stimulus. **d**) Examples of orofacial motion capture from an animal. **e**) Change in absolute motion energy plotted relative to lickspout arrival and compared to baseline (1-0.1 s before stimulus onset). **f**) Change in absolute motion energy relative to stimulus onset and compared to baseline (1 - 0.1 s before stimulus onset). Inset: zoom of first 0.6 s, visualizing the fast onset of the rise in motion energy in conditions with an auditory stimulus. **g**) Z-scored motion energy relative to the pre-stimulus baseline period for stimulus locked and lickspout-locked conditions. **h**) Z-scored motion energy in response to stimulation with an auditory component. All data show mean (± SEM). *p<0.05; p<0.01; p<0.001; n.s. = not significant.

### Responses of V1 neurons become less selective to stimulus type throughout the trial

Introduction of a movable lickspout induced an increase in orofacial motion after lickspout onset (Fig. 1e, repeated measures ANOVA, effect of time: F(2,8)= 277.49, p=4.08e-08; effect of modality F(3,12)=0.50, p=0.69; interaction effect: F(6,24) = 0.39, p=0.88). During the stimulus-only period we observed a small increase in videoME during auditory or multisensory, but not visual or no-stimulus trials (Fig. 1f). A strong increase in motion energy was found in the lickspout-locked period compared to baseline (Fig. 1g, Bonferroni corrected paired t-test: Auditory: 2.19±0.46, p=0.0014; Visual: 1.93±0.78, p=0.016; Multisensory: 1.93±0.46, p=0.0022; No Stimulus: 1.85±0.22, p=0.00015). Furthermore, stimulus saliency may affect stimulus-induced orofacial motion. Indeed, auditory intensity, but not visual contrast, was correlated with the increase in orofacial motion energy after stimulus presentation (Fig. 1h, Fig. S1, effect of auditory intensity: F(3,12)= 3.93, p=0.036; effect of modality (A or M): F(1,4)= 0.12, p=0.75; interaction effect: F(3,12)= 1.55, p=0.25; effect of visual contrast: F(13,12)= 2.48, p=0.11) in a subset of sessions (Fig. S1).

To investigate neural correlates of stimulus perception and orofacial movement, we recorded left V1 with laminar silicon probes during task performance (Fig. 2a). We recorded 193 units across 15 sessions in five mice. Single units showed stimulus- and lickspout-locked responses (example cells in Fig. 2a). The number of responsive units increased as the trial progressed and became less selective to stimulus type (Fig. 2b). If single units significantly increased their firing in the stimulus-only period compared to baseline (1 to 0.1 s before stimulus onset) we labelled them as visually (i.e. responding to V or V+M, but not A trials, N=72) or auditory selective (i.e. responding to A or A+M, but not V trials, N=11). We performed a generalized linear regression on counts of these cell types across epochs. The GLM coefficients showed that the fraction of visually selective units that increased their firing during V and M trials decreased across the trial (Fig 2c, binomial GLM coefficients, stim-only vs. lickspout-arrival: -0.73±0.23, p=0.00049; lickspout-arrival vs. response/reward: -1.17±0.25, p=2.28e-06), whereas the fraction of units that significantly increased their firing rate during no-stimulus trials increased (Fig 2c, binomial GLM coefficients, stim-only vs. lickspout-arrival: 2.47±0.29, p<2e-16; lickspout-arrival vs. response/reward: 2.91±0.29, p<2e-16). The fraction of units that significantly increased their firing only during A and M trials (i.e., auditory selective units) increased slightly during the lickspout-arrival epoch (Fig. 2c, binomial GLM coefficients, stim-only vs lickspout-arrival: 0.81±0.38, p=0.035; lickspout-arrival vs reward/response: 0.65±0.39, p=0.096). Taken together, the responses of single units became less selective to stimulus type throughout the trial.

**Figure 2.**
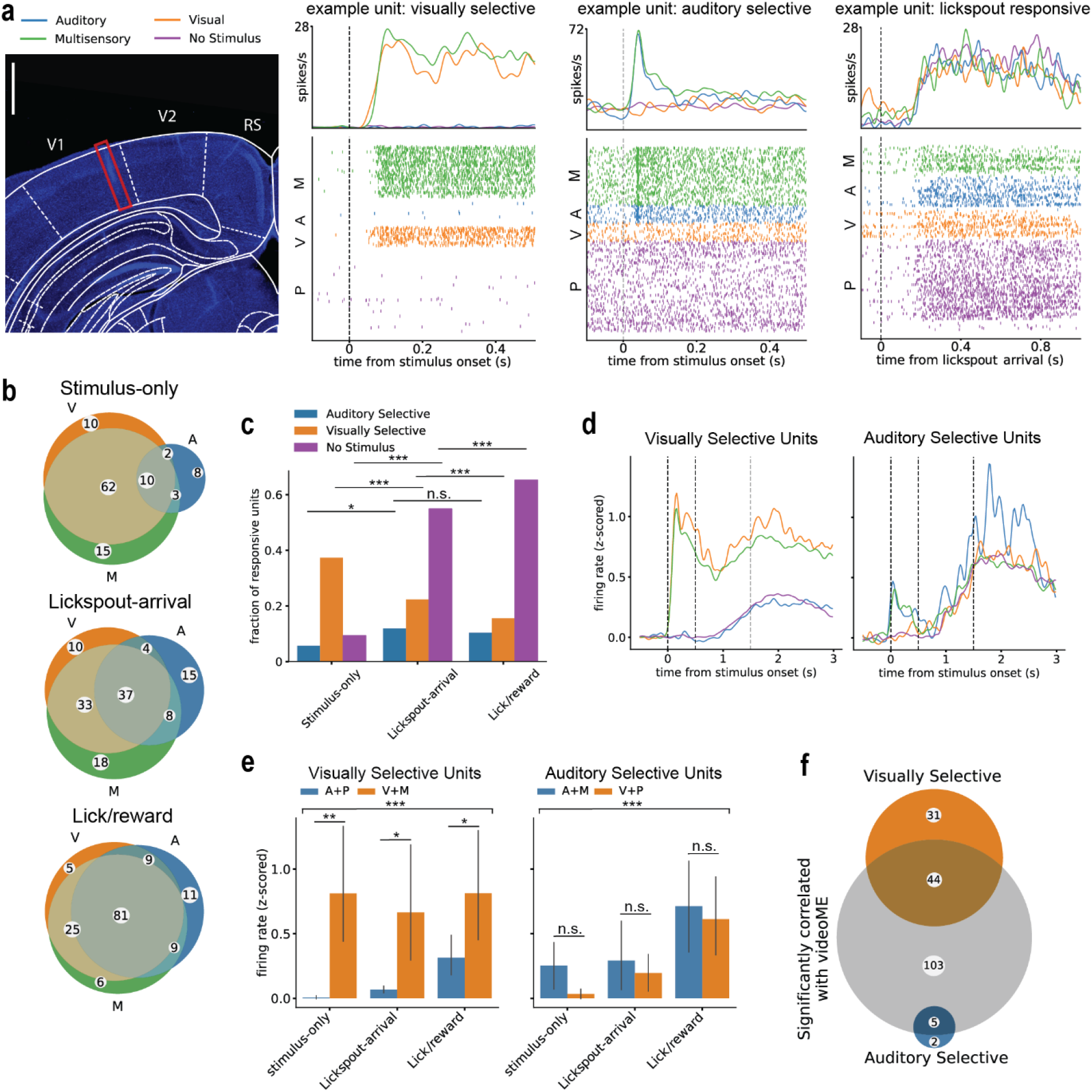
Correlation of visually and auditory selective V1 units with motion across the entire trial. **a**) Example of inserted probe in mouse V1 (left) and example visually selective, auditory selective, and lickdetector responsive neurons, respectively (right). Calibration bar is 500 um. **b**) Stimulus selectivity of V1 neurons in the stimulus-only, lickspout arrival, and lick/reward epochs. **c**) Fractions of cells significantly responsive to conditions with auditory stimuli, visual stimuli or no stimulus for the three behavioral epochs: stimulus-only (0 – 0.5 s), lickspout-arrival (0.5 – 1.5 s), lick/reward (1,5 – 4 s). **d**) Z-scored average firing rate responses of visually and auditory selective units relative to stimulus onset. Dashed lines on 0.5 and 1.5 s indicate the lickspout-arrival epoch, coinciding with an increase in neuronal and behavioral activity. For color legend, see (a). Visual and auditory selectivity were determined in the stimulus-only period (0 – 0.5 s). **e**) Difference in Z-scored firing rates, along the three epochs, for non-corresponding and corresponding stimuli: i.e. for visually selective cells (left, N=72), firing rates in auditory and probe trials (A + P; non-corresponding to sensory selectivity) versus visual and multisensory trials (V + M; corresponding); for auditory selective units (N=11), firing rates to auditory and multisensory trials (A + M; corresponding) versus visual and probe trials (V + P; non-corresponding). **f**) Cells showing a significant correlation between firing activity and motion energy (videoME), computed across the entire trial. Orange and blue show numbers of visually (N=75 in total) and auditory selective cells (N=7), respectively. The grey circle indicates the number of cells of which the firing rate significantly correlated with videoME. The two intersections show selective cells significantly correlating with videoME. All data show mean (± SEM). *p<0.05; p<0.01; p<0.001; n.s. = not significant.

Next, we focused on units that were selective in the first epoch, the stimulus-only period. The visually selective population showed a stimulus-locked response during visual and multisensory trials, which was followed by a second bump of late activity during the lickspout-arrival and response/reward period (Fig. 2d,e, main of epoch (time period): F(2,136)= 10.09, p=8.21e-05; main effect of modality: F(1,68)= 9.46, p=0.0030; interaction effect: F(2,136)= 12.43, p=1.11e-05; post-hoc Bonferroni corrected F-test A+P vs V+M trials: stimulus-only: p=0.0025; lickspout-arrival: p=0.026; response/reward: p=0.015). During this period of late activity, the z-scored firing rates of visually-selective neurons (N=72) also increased during auditory and no-stimulus trials (stimulus-only: 0.01±0.05; lickspout-arrival: 0.07±0.11; response/reward: 0.31±0.67), but these were lower than the z-scored activity during visual and multisensory trials (Fig. 2e, stimulus-only: 0.81±1.91; lickspout-arrival: 0.67±0.88; response/reward: 0.81±1.89). The auditory selective population showed similar patterns, except that there was no difference in late activity between auditory, visual, multisensory or no stimulus trials (Fig. 2d,e, main effect of epoch: F(2,10)= 15.64, p=0.00083; no effect of modality: F(1,5)= 2.72, p=0.16; no interaction effect: F(2,10)= 1.72, p=0.23). Thus, these results show stimulus-selective responses during the stimulus-only period, but this selectivity largely disappears later in the trial, once the mouse could make its response to obtain reward.

We next investigated whether the activity of stimulus-type selective and nonselective units correlated with orofacial motion before and after lickspout onset (Fig. 2f). Of the recorded units, over all trials and for the whole duration of the trial, 53.4% (103/193) were significantly correlated with orofacial motion (defined by their Pearson correlation, p<0.05, Bonferroni-corrected circular shuffle test). Of the visually selective cells, 58.7% (44/75) correlated to orofacial motion. This fraction was 71.4% (5/7) for the auditory selective units (Fig. 2f). This indicates that on a population level, across the entire trial, visual as well as auditory information is affected by orofacial motion. However, this leaves open the question how neural activity is correlated with orofacial motion in different epochs of a trial.

### A subpopulation of V1 neurons shows minor motor correlates after visual, but not auditory stimulus onset

Thus, to further examine whether stimulus and motor information are separately coded, we trained a Random Forest model to predict single-trial firing rates over time and for distinct trial epochs (Fig. 3a,b). This model included stimulus-, motor-, lickspout- and trial-specific predictors (see Methods). Additionally, we trained two more models, one excluding stimulus-specific (stimulus presence and intensity/contrast) and one excluding motor-specific (orofacial motion and licking) predictors. These showed that the stimulus contribution to the variance in firing rate decreases over trial time, whereas the contribution of motor variables increases (Fig. 3c, main effect of epoch: F(2,384)=7.77, p<0.001; no main effect of predictor set: F(1,192)=2.23, p=0.14; interaction effect: F(2,384)=65.62, p<0.001). The number of units significantly encoding stimulus information remained stable throughout the trial (Fig. 3d, binomial GLM coefficients, stim-only vs. lickspout-arrival: -0.08±0.20, p=0.693; lickspout-arrival vs. reward/response: 0.11±0.19, p=0.566). In the stimulus-only period, 26% of the units (Fig. 3d, 50/193; binomial test, p<0.001) significantly encoded motor predictors, which throughout the trial significantly increased in number (Fig. 3d, binomial GLM coefficients, stim-only vs. lickspout-arrival: 0.72±0.19, p<0.001; lickspout-arrival vs. reward/response: 1.09±0.19, p<0.001). Thus, as the trial progressed, more units came to encode motor information and more variance in firing rate could be explained by orofacial motion. At a population level, the stimulus-only period, before mice can make a behavioral response, was generally not free of motor confounds.

**Figure 3.**
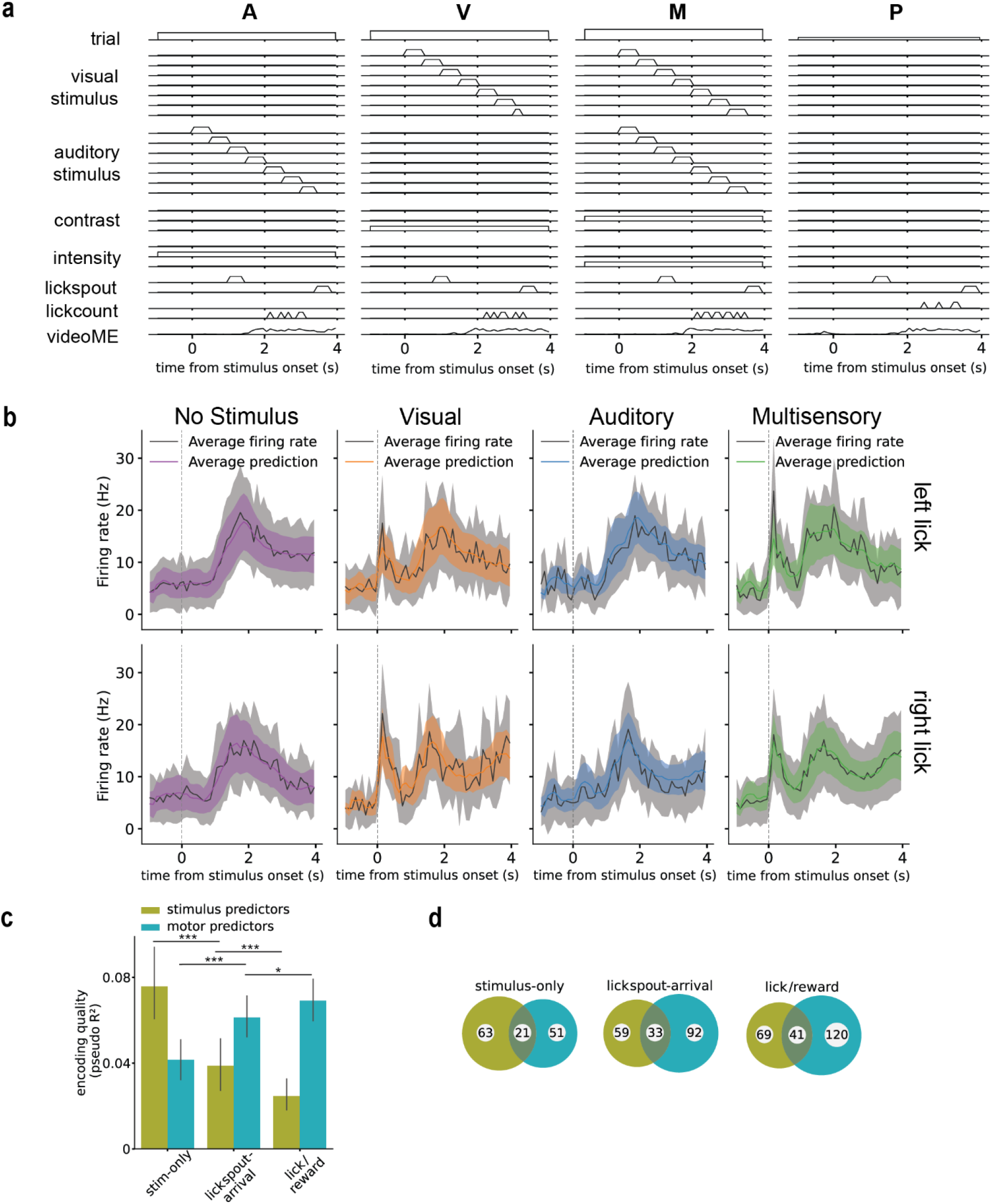
Encoding of stimulus and motor information by V1 neurons. **a**) Illustration of the parameters used to train the Random Forest model. Stimulus predictors included stimulus presence and contrast/intensity. Motor predictors included orofacial motion and licking parameters. The lickspout predictors indicate the movement of the lickspout towards the mouse and away from the mouse, and thus the lick/response epoch. **b**) Predicted firing rates from the Random Forest model (color) and the actual firing rates (grey) of an example cell, for all four stimulus conditions (columns) and left and right licking responses (rows). Data shows mean (± 0.5 SD) **c**) Encoding quality of Random Forest models trained with either stimuli predictors or motor predictors, over the three behavioral epochs: stimulus-only (0 – 0.5 s), lickspout-arrival (0.5 – 1.5 s), lick/reward (1,5 – 4 s). **d**) Numbers of units for which stimulus (olive) and/or motor predictors (cyan) were significant (as defined by a significant difference between encoding models with and/or without said predictors) over the three behavioral epochs. All data show mean (± SEM), except where stated otherwise. *p<0.05; p<0.01; p<0.001; n.s. = not significant.

We wondered whether this finding holds true for all subpopulations of neurons. During visual and multisensory trials, visually selective units (N=69) showed a stimulus component in the stimulus-only window significantly higher than zero (Fig. 4a, visual trials: t(68)=8.01, p<0.001; multisensory trials: t(68)=9.37, p<0.001; Bonferroni corrected). This stimulus component decreased as mice produced their instrumental response in visual and multisensory trials (Fig. 4a, main effect of epoch: F(2,136)=57.05, p<0.001; no main effect of modality: F(1,68)=0.46, p=0.50; interaction effect: F(2,136)=4.25, p=0.02). This contribution of visual stimulus information increased with stimulus contrast (Fig. 4c, main effect of epoch: F(2,136)=56.76, p<0.001; main effect of contrast: F(2,136)=54.77, p<0.001; interaction effect: F(4,272)=22.25, p<0.001). Contrarily, there was no significant motor contribution in the stimulus-only window, before lickspout movement (Fig. 4a, visual trials: t(68)=2.09, p=0.48; multisensory trials: t(68)=2.96, p=0.05; Bonferroni corrected), but this component increased as the trial progressed (Fig. 4a, main effect of epoch: F(2,136)=10.90, p<0.001; no main effect of modality F(1,68)=0.08, p=0.77; no interaction effect: F(2,136)=0.45, p=0.64). This suggests that motor confounds in this visually selective subpopulation remain low in this initial window during visual stimulus presentation, before mice could make a behavioral response. Nevertheless, a small amount of spontaneous movements in the stimulus-only period during no-stimulus trials remained (Fig. S2a, all units: t(192)=7.56, p<0.001; visually selective units: t(68)=5.76, p<0.001; Bonferroni corrected), which was not significantly different from the general population (Fig. S2a, GLM coefficient visual selectivity: 0.001±0.007, p=0.86). These conclusions do not hold true for auditory selective units (N=7; Fig. S3a). Those units showed a consistently strong motor component (Fig. S3a, no main effect of epoch F(2,12)=1.63, p=0.24; no main effect of modality F(1,6)=1.00, p=0.36; no interaction effect F(2,12)=0.86, p=0.44) and modest to no stimulus component (Fig. S3a, no main effect of epoch F(2,12)=2.60, p=0.12; no main effect of modality F(1,6)=1.54, p=0.26; no interaction effect F(2,12)=0.53, p=0.60) across all time windows. Relevant post-hoc tests showed no significant stimulus component in the stimulus-only period (Fig. S3a, auditory trials: t(6)=0.83, p=0.44; multisensory trials t(6)=1.37, p=0.22; Bonferroni corrected). Additionally, only a few auditory selective neurons were found to significantly encode either stimulus (N=1) or motor (N=4) components or both (N=1) in the stimulus-only period (Fig. S3a). Together, these results indicate that neural V1 responses after auditory stimulus presentation are mainly explained by the orofacial movement induced by the auditory stimulus. These results show that, when applying the delayed-reward method, motor-related activity does not affect a subset of visually selective V1 neurons, whereas auditory responses are confounded by motor-related activity, even in the stimulus-only period of the trial.

**Figure 4.**
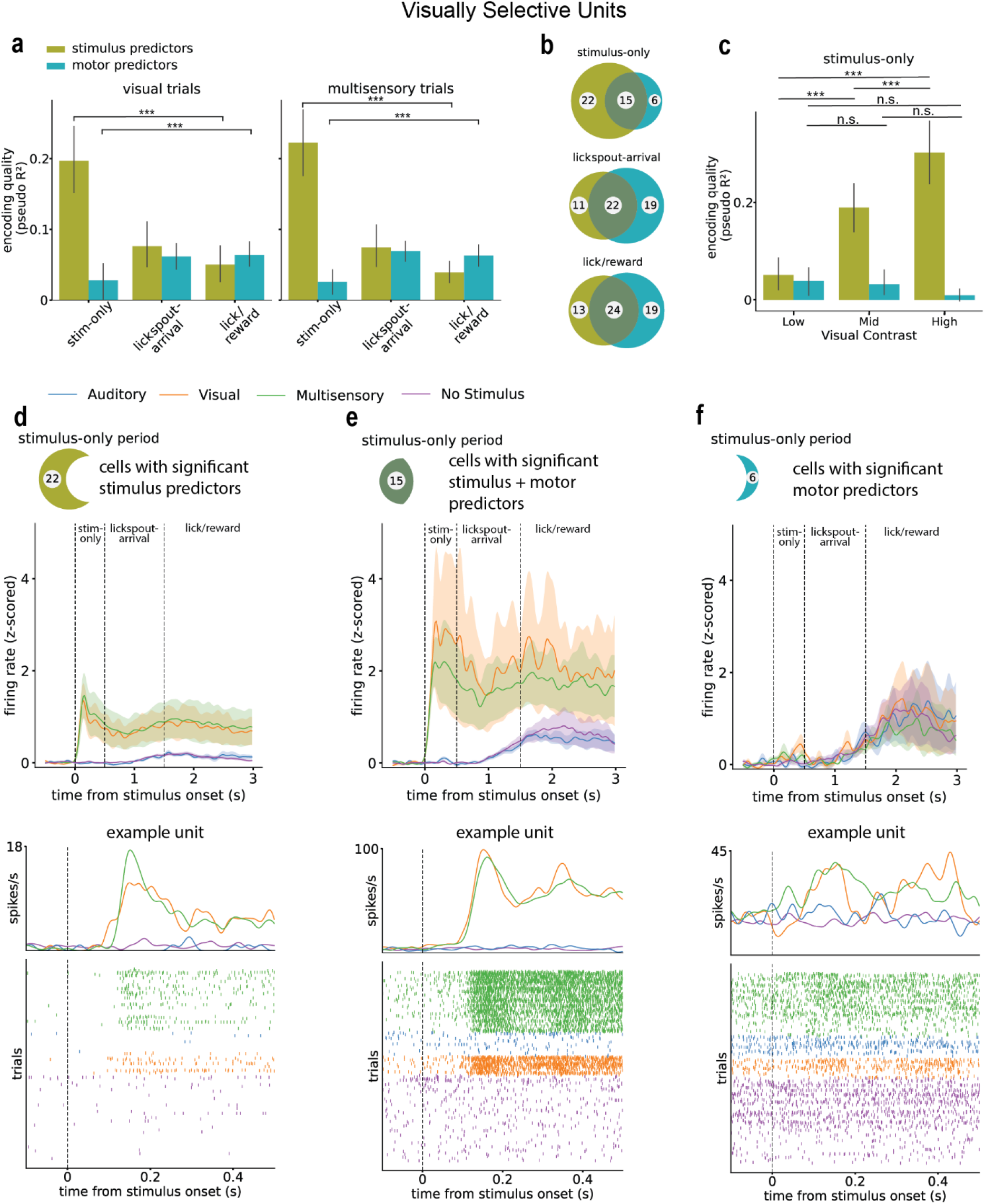
Encoding of stimulus and motor predictors in visually selective V1 neurons. **a**) Encoding quality of Random Forest models trained with either stimulus predictors or motor predictors, over the three behavioral epochs: stimulus-only (0 – 0.5 s), lickspout-arrival (0.5 – 1.5 s), lick/reward (1,5 – 4 s) in visual (left) and multisensory (right) trials. **b**) Numbers of units for which stimulus (olive) and/or motor predictors (cyan) were significant (as defined by a significant difference between encoding models with and/or without said predictors) over the three behavioral epochs. **c**) Encoding quality of stimulus (olive) and motor (cyan) predictors across visual stimulus contrast in the stimulus-only period (0 – 0.5 s). **d**) Z-scored stimulus-locked firing rate responses of cells with significant stimulus predictors in the stimulus-only period (0 – 0.5 s) for all trial types (top) and an example neuron (bottom). Crescent shapes refer to the venn-diagram in figure b. **e**) Z-scored stimulus-locked firing rate responses of cells with significant stimulus and motor predictors in the stimulus-only period (0 – 0.5 s) for all trial types (top) and an example neuron (bottom). **f**) Z-scored stimulus-locked firing rate responses of cells with significant motor predictors in the stimulus-only period (0 – 0.5 s) for all trial types (top) and an example neuron (bottom). All data show mean (± SEM). *p<0.05; p<0.01; p<0.001; n.s. = not significant.

## Discussion

We trained mice on an audiovisual detection task and forced a delay in their behavioral response by postponing placement of the reward apparatus in reach of the mouse until well after stimulus onset. This yielded a stimulus-only time window where the animals were unable to lick at the dual lickspout, and we hypothesized that orofacial motion during this window would be minor. We studied whether this approach could prevent major motor correlates from arising in V1 neuronal responses in the time window directly following stimulus onset. After stimulus onset, but before lickspout arrival, mice indeed showed low spontaneous orofacial motion. Arrival of the lickspout induced profuse orofacial movements and licking, which was correlated to spiking activity in V1. Additionally, salient auditory, but not visual, stimuli evoked notable orofacial movement. Accordingly, auditory responses in V1 were largely driven by orofacial motion. However, a subpopulation of neurons which responded selectively to visual stimulation carried information primarily about the visual stimulus before a behavioral response could be made (Fig. 4). In contrast, the activity of auditory selective V1 units correlated strongly with orofacial movement (Fig. S3). Thus, by using a movable dual lickspout in a visual detection task, motor confounds in spiking activity remain very limited in a subset of visually responsive neurons in the stimulus-only period, even though such confounds should still be controlled for.

### Differences in motor confounds between visually and auditory-selective V1 neurons

Spontaneous movements during visual stimulation are correlated with brain-wide activity^4^. Visual stimulation rarely elicits such a strong orofacial response as auditory stimulation does, and to the extent such as response is evoked, it does not explain much of the visually related neuronal response in auditory cortex^14^. Here, we physically delayed the arrival of the reward apparatus; despite this forced delay, a high percentage of all recorded neurons was significantly correlated with orofacial movement, even if videomotion energy was relatively low. However, a subpopulation which selectively responded to visual stimulation primarily carried information on the visual stimulus, as evident from a high encoding quality in the stimulus-only period, but not on orofacial motor activity, before a licking response could be made (Fig. 4a,c,d and 2d). Contrarily, auditory responsive V1 units mostly conveyed information on orofacial motion (Fig. S3). These results indicate that, with a forced delay in introducing the reward apparatus, motor correlates are limited in a subset of units. Interestingly, the cortex-wide activity ignited in preparation of a behavioral response^2,4,15^ thus appears not to be a generalized wave of activity affecting all V1 cells equally, but only affects a subset of them. The function of such motor correlates remains to be elucidated, but may well relate to the importance of motor efference copy and/or proprioceptive reafferency from active musculature in the integration with retinal information, in order to stabilize the subject’s world view in the face of ongoing body movement^35,36^. The fact that only a subset of visually responsive neurons shows motor correlates appears to align with the highly structured and selective computational architecture that would be needed to perform these functions, compared to a blanket modulation of all visual responses. The movable lickspout may well serve as a Pavlovian cue predicting reward. Bringing this cue into proximity of the mouse may elicit an enhanced instrumental response after stimulus onset through Pavlovian-Instrumental Transfer (PIT)^17,37,38^. We observed that orofacial movement was strongly enhanced upon lickspout arrival (Fig. 1e), which is likely coupled to enhanced motivational arousal. However, also the visual and auditory stimuli may act as Pavlovian cues, which may partly explain the increase in orofacial movement after auditory stimulus onset (Fig. 1f). However, cue-induced Pavlovian conditioning during behavioral training would have to be demonstrated in a separate experiment. Apart from differences in subjective saliency, it remains unclear, however, why visual cues were less effective than auditory cues in evoking orofacial motion. Modulation of behavior by Pavlovian conditioning is thought to be mediated, at least in part, by affective cortico-basal ganglia loops and the associated dopaminergic cells in the ventral tegmentum^17,25,39–41^. It would be interesting to directly investigate whether the introduction of a delayed behavioral response in animals trained without a delay also modulates V1 responses to visual stimulation in a similar fashion. Because we introduced a random delay (between 0.5-1.5 s) in lickspout arrival, the timing of licking and reward acquisition becomes less predictable than with a fixed interval. Whether this temporal variation is also reflected in anticipatory activity to conditioned stimuli, as shown before^19^, remains an open question.

### Sound evokes orofacial motion and neural activity in V1

In line with the prior literature, we found that auditory stimulation evoked orofacial movements and responses in V1^2,3,14,15^. This was most clearly reflected in auditory responsive cells, in which orofacial movement accounts for most of the explained variance in neural activity (Fig. S3). In V1, auditory inputs arrive early on (∼ 27 ms post-stimulus onset^9^), before the onset of motor confounds^3^. Here, we looked at different temporal epochs, but did not find auditory V1 responses free of motor confounds. This is likely due to the fact that we adopted a time window for analysis of 0.5 s post-stimulus onset (i.e., the stimulus-only period), during which orofacial motor correlates will already unfold. In addition, these results might be skewed due to the low number of auditory responsive units in our dataset. This study focused on V1 data, however, motor-related signals have been found cortex-wide^4,42^. For example, in auditory cortex units start firing before the onset of orofacial movements associated with sound representation and reward^43^, which could represent the learned link between the auditory stimulus and expected reward^44^. This illustrates that sensory cortical areas, such as V1 and auditory cortex, do not only process and relay sensory information^35^. Motor signals found in V1 may play a functional role as efference copy signals, which convey predictions on what the visual input will be, given the motor movement^36,45^. Thus, motor signals do not only confound neural data, but likely also serve a function in natural behaviors^42^.

### Technical limitations

One of the limitations of our study is that we did not include a control experiment that compared the same task without a delayed response window. Hence, we refrain from making a claim to the effect that delayed insertion would exert a causal influence on diminishing orofacial motion. We have previously shown that auditory stimulation without delayed introduction of the reward apparatus also elicits orofacial movement correlates in V1^2,3^, but one cannot use the current findings to estimate whether a delayed introduction reduces this type of orofacial motion. To control for spontaneous movement, we included ‘no stimulus’ (probe) trials, in which mice are not presented a stimulus before lickspout arrival. This excludes any motion evoked by stimulus presentation. We found that spontaneous motion indeed contributed to neural variance (Fig. S2a). Here, the motion contribution to neural variance of the entire population of V1 units did not differ from that observed for visually selective units (Fig. S2a). This indicates that, in this subpopulation of units, visual stimulation does not evoke orofacial movement-related activity in addition to spontaneous activity, which agrees with our finding that visual stimulation evoked very little videomotion energy after stimulus onset to begin with. The current study allows the experimenter to control for spontaneous movements before the behavioral response, but it does not abolish these completely. The later onset of the lickspout provides an opportunity to better demarcate motor-related and sensory information in the primary visual cortex. However, controls for motor confounds remain necessary.

### Conclusion

The main finding of our study is the identification of a subset of visually responsive V1 neurons that did not show correlations with orofacial movement in the trial period where only the stimulus was present. In contrast, sound-responsive V1 cells generally do express motor-related activity shortly after stimulus onset. Thus, even with delayed rewards, it remains necessary to control for motor confounds in neural data of head-fixed awake mice. Stimulus presentation can induce orofacial motion and spontaneous movements remain. Nonetheless, with this method the impact of these confounds can be minor, at least in a subset of cells.

## Acknowledgements

This work was supported by the Netherlands Organization for Scientific Research-VICI Grant 918.46.609 (to CMAP), the EU FP7-ICT grant 270108 (to CMAP), the European Union’s Horizon 2020 Framework Programme for Research and Innovation under the Specific Grant Agreement No. 945539 (Human Brain Project SGA3 to CMAP), and the NWO Crossover project INTENSE to U.O. and C.M.A.P.

---

**Figure S1.**
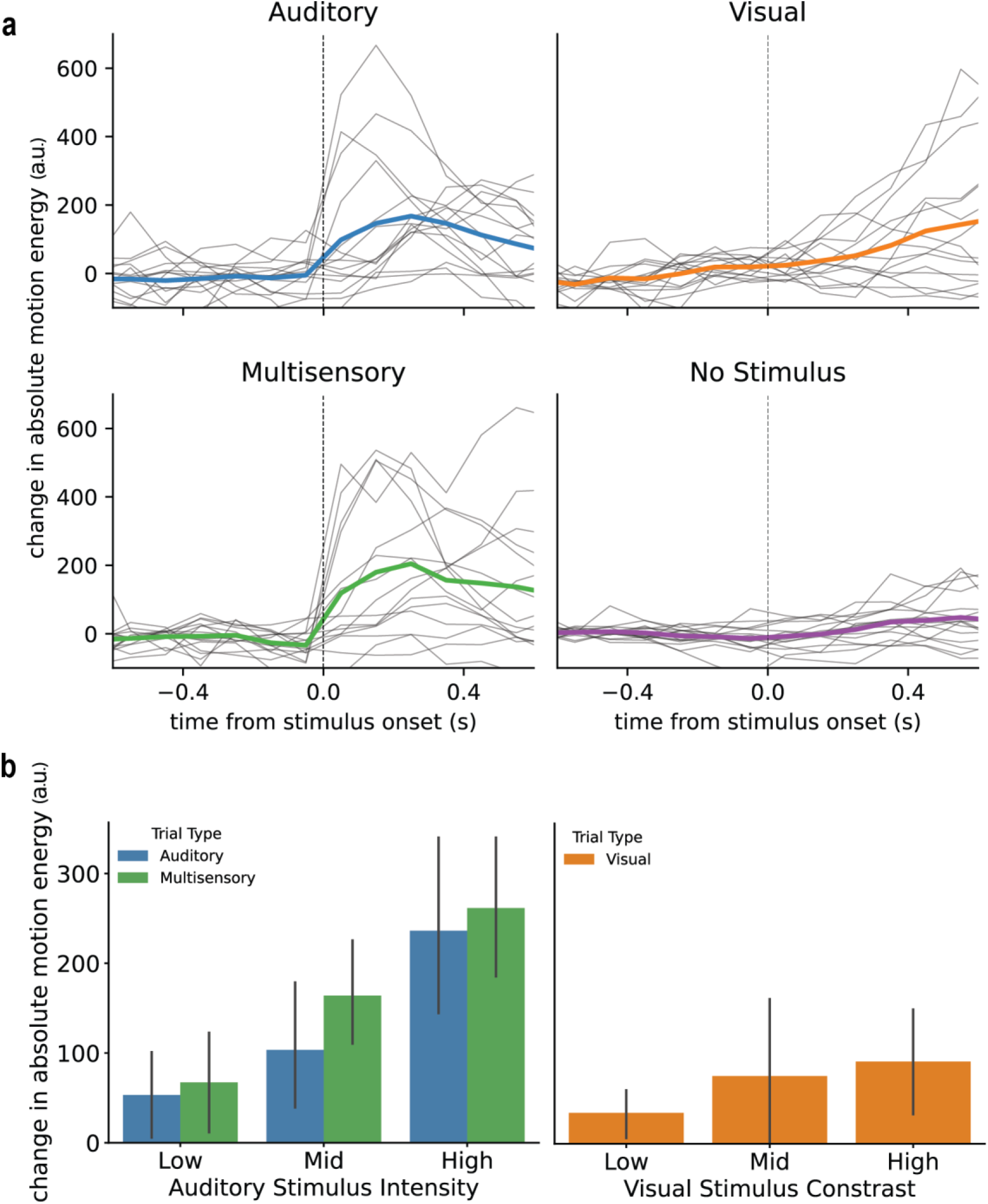
Motion energy change due to different types of stimulus. **a**) Average motion energy changes (grey lines: individual sessions, thick line: average) after stimulus onset compared to baseline for auditory, visual, multisensory and probe conditions. **b**) Changes in motion energy mean (±SEM) for different auditory intensities (left panel) and visual contrasts (right panel) in the stimulus-only window. Multisensory trials grouped by visual contrasts are not plotted since those include auditory stimuli of varying intensities. The stimulus intensity and contrast are defined per individual based on its detection threshold (see Methods).

**Figure S2.**
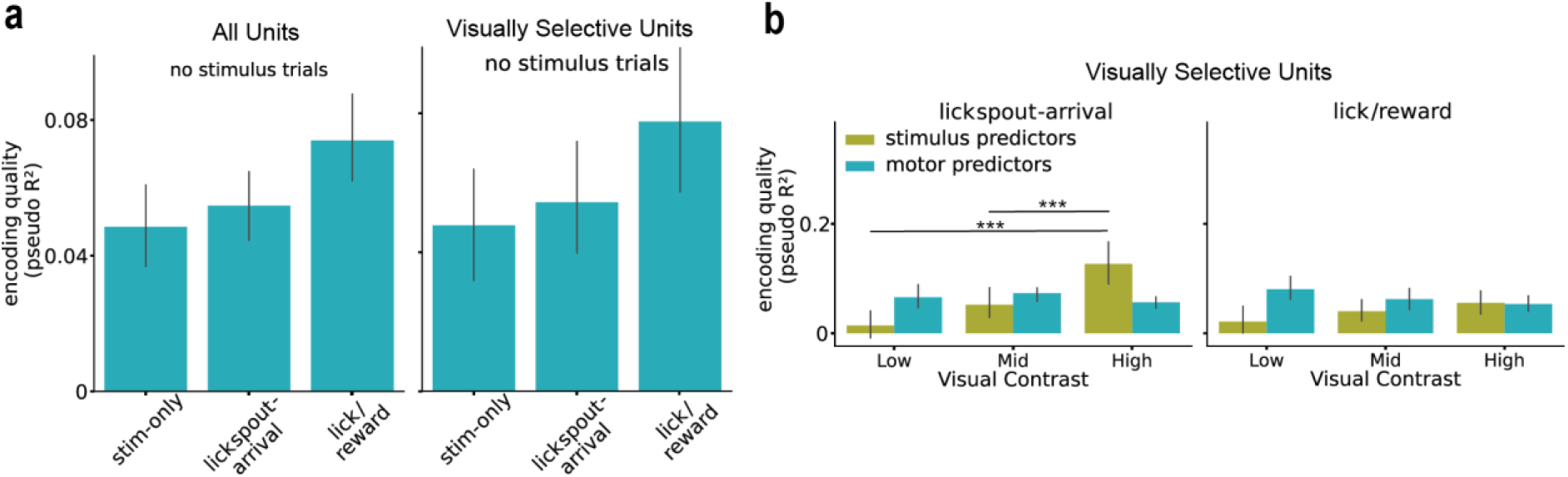
Encoding of firing rates in visually selective V1 units. **a**) Encoding quality of motor predictors across behavioral epochs during the no-stimulus (probe) trials in all units (left) and visual selective units (right). **b**) Encoding quality across visual stimulus contrasts for visually selective V1 units with either stimulus or motor predictors in the encoding model during the lickspout-arrival (1 – 1.5 s; left) and lick/reward period (1.5 – 4 s; right).

**Figure S3.**
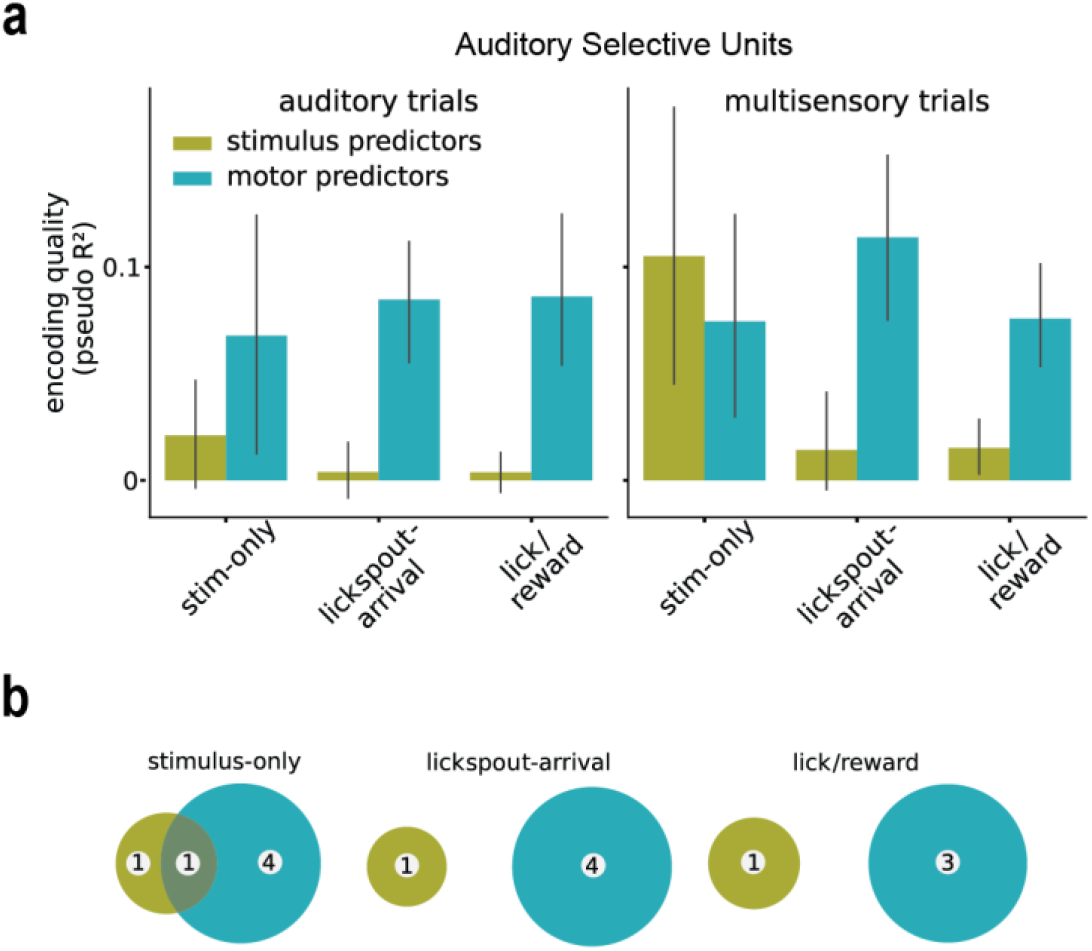
Encoding of firing rates in auditory selective V1 neurons. **a**) Encoding quality of stimulus and motor predictors across behavioral epochs in auditory (left) and multisensory (right) trials. **b**) Numbers of units for which stimulus (olive) and/or motor predictors (cyan) were significant (as defined by a significant difference between encoding models with and/or without said predictors) over the three behavioral epochs.

